# Sleep-dependent modulation of metabolic rate in *Drosophila*

**DOI:** 10.1101/124198

**Authors:** Bethany A. Stahl, Melissa E. Slocumb, Hersh Chaitin, Justin R. DiAngelo, Alex C. Keene

## Abstract

Dysregulation of sleep is associated with metabolic diseases, and metabolic rate is acutely regulated by sleep-wake behavior. In humans and rodent models, sleep loss is associated with obesity, reduced metabolic rate, and negative energy balance, yet little is known about the neural mechanisms governing interactions between sleep and metabolism. We have developed a system to simultaneously measure sleep and metabolic rate in individual *Drosophila*, allowing for interrogation of neural systems governing interactions between sleep and metabolic rate. Like mammals, metabolic rate in flies is reduced during sleep and increased during sleep deprivation suggesting sleep-dependent regulation of metabolic rate is conserved across phyla. The reduction of metabolic rate during sleep is not simply a consequence of inactivity because metabolic rate is reduced ∼30 minutes following the onset of sleep, raising the possibility that CO_2_ production provides a metric to distinguish different sleep states in the fruit fly. To examine the relationship between sleep and metabolism, we determined basal and sleep-dependent changes in metabolic rate is reduced in starved flies, suggesting that starvation inhibits normal sleep-associated effects on metabolic rate. Further, *translin* mutant flies that fail to suppress sleep during starvation demonstrate a lower basal metabolic rate, but this rate was further reduced in response to starvation, revealing that regulation of starvation-induced changes in metabolic rate and sleep duration are genetically distinct. Therefore, this system provides the unique ability to simultaneously measure sleep and oxidative metabolism, providing novel insight into the physiological changes associated with sleep and wakefulness in the fruit fly.

**Significance statement:** Metabolic disorders are associated with sleep disturbances, yet our understanding of the mechanisms underlying interactions between sleep and metabolism remain limited. Here, we describe a novel system to simultaneously record sleep and metabolic rate in single *Drosophila*. Our findings reveal that uninterrupted sleep bouts of 30 minutes or greater are associated with a reduction in metabolic rate providing a physiological readout of sleep. Use of this system, combined with existing genetic tools in *Drosophila*, will facilitate identification of novel sleep genes and neurons, with implications for understanding the relationship between sleep loss and metabolic disease.

## Introduction

Dysregulation of sleep is strongly linked to metabolism-related pathologies and reciprocal interactions between sleep and metabolism suggest these processes are integrated at the cellular and molecular levels^1,2^. In mammals, metabolic rate is reduced during sleep raising the possibility that sleep provides a mechanism of energy conservation or partitioning^3^. While a reduction in metabolic rate and energy expenditure during sleep has been documented in mammalian and avian species^4,5^, little is known about the genetic and neural mechanisms governing the effects of sleep on metabolic rate. The fruit fly, *Drosophila melanogaster*, displays all the behavioral characteristics of sleep and provides a powerful system for genetic investigation of interactions between sleep and diverse physiological processes^2,6,7^. Here, we describe a novel single-fly respirometry assay in the fruit fly, designed to simultaneously measure sleep and whole-body metabolic rate that allows for genetic interrogation of the mechanisms regulating interactions between these processes.

Sleep is characterized by physiological changes in brain activity or through the behavioral correlates that accompany these changes^8^. Flies, like mammals, display distinct electrophysiological patterns that correlate with wake and rest^9,10^. Additionally, flies display all the behavioral hallmarks of sleep including extended periods of behavioral quiescence, rebound following deprivation, increased arousal threshold, and species-specific posture^6,7^. Sleep in *Drosophila* is typically defined by five minutes of behavioral quiescence, because this correlates with other behavioral characteristics used to define sleep^7^. While these behavioral metrics of sleep have been studied extensively, significantly less is known about physiological changes associated with sleep in flies.

In rodents and humans, metabolic rate is elevated in response to sleep-deprivation and reduced during sleep, supporting the notion that metabolic processes are acutely regulated by sleep state^11–13^. In flies and other small insects, stop-flow respirometry can be used to monitor CO_2_ production, a by-product of oxidative metabolism and a proxy for metabolic rate^14^. Here, we describe a system to simultaneously measure sleep and metabolic rate in individual fruit flies. Our findings reveal that metabolic rate is reduced when flies sleep, and uninterrupted sleep bouts of ∼30 minutes or greater are associated with an additional reduction in metabolic rate, indicating that flies exhibit sleep stages that are physiologically distinct. Further, we find that starvation inhibits sleep-associated reductions in metabolic rate, suggesting feeding state influence physiological changes associated with sleep. These findings suggest that sleep-dependent reductions in metabolic rate previously observed in mammals are conserved in the fruit fly, and further support the notion that sleep provides a mechanism for energy conservation.

## Methods

### *Drosophila* maintenance and Fly Stocks

Flies were grown and maintained on standard food (Bloomington Recipe, Genesee Scientific). Flies were maintained in incubators (Powers Scientific; Dros52) at 25°C on a 12:12 LD cycle, with humidity set to 55-65%. The wild-type line used in this manuscript is the *w*^1118^ fly strain (Bloomington Stock #5905). The trsn^null^ allele is an excision of the *trsn*^EY06981^ locus derived from mobilizing the EPgy2 insertion^15^. This allele removes the entire coding region of the gene and represents a null mutation that has been outcrossed to the *w*^1118^ background and has previously been described as Δ*trsn*^15^. Unless noted in the figures, all experiments are performed in 3-5 day old mated female flies.

### Measurement of metabolic rate and locomotor activity

Metabolic rate was measured at 25°C through indirect calorimetry, measuring the CO_2_ production of individual flies with a Li-7000 CO_2_ analyzer (LI-COR), which was calibrated with pure CO_2_ prior to each run. A stop-flow, push-through respirometry setup was constructed using Sable Systems equipment (Sable Systems International). The experimental setup included sampling CO2 from an empty chamber to assess baseline levels, alongside five behavioral chambers, each measuring CO_2_ production of a single fly. The weight flies used for analysis were not taken into account because body size and energies stores are not perturbed in *trsn*^*null*^ flies and do not vary significantly in *w*^1118^ flies. Further, previous work using a comparable system suggests weight will have little effect on CO_2_ measurements unless there is an excess of >50% differences in size between individuals^14,16^. To measure CO_2_ output, air was flushed from each chamber for 50 seconds providing a readout of CO_2_ accumulation over a 5-minute period. This 5-minute interval allows the coordinate and simultaneous activity-based assessment of sleep. The first 20 minutes of recordings were not included in analyses because this time was necessary to purge the system of ambient air and residual CO_2_ from the closed system. Dehumidified, CO_2_ free air was pumped through a mass flow control valve (Side-Trak 840 Series; Sierra Instruments, Inc.) to maintain the experimental flow rate of 100mL/min. The air was then passed through water-permeable Nafion tubing (Perma Pure, LLC, Lakewood, NJ USA, #TT-070) immersed in a reservoir containing deionized H_2_O to re-humidify the air prior to reaching the behavioral chambers. Non-permeable Bev-A line tubing (United States Plastic Corp., Lima, OH USA, #56280) was used throughout the rest of the system.

Experiments were conducted by placing single flies in 70mm × 20mm glass tubes that fit a custom built *Drosophila* Locomotor Activity Monitor (Trikinetics, Waltham, MA) with three sets of infrared beams for activity detection. The monitor was connected to a computer to record beam breaks every minute for each animal using standard DAMS activity software (Trikinetics, Waltham, MA) as previously described^17^. These data were used to calculate sleep information by extracting immobility bouts of 5 minutes using a custom generated python program. The total activity from all three beams was summed for each time point in order to determine overall activity. Video recordings for analysis of feeding activity were acquired using a handheld USB Digital microscope (Vivida, 2MP #eheV1-USBpro) camera at 12 fps VirtualDub software (v.1.10.4). Each 60-minute video recording occurred between ZT01-04 to prevent circadian differences in sleep, feeding and metabolic rate. During video recording, flies were simultaneously assayed for activity and metabolic rate, with the stop flow set to collect CO_2_ output every 2 minutes. Videos were manually scored for feeding activity in corresponding 2-minute intervals as a “feeding” or “non-feeding” bin.

Flies were briefly anesthetized using CO_2_ for sorting at least 24hrs prior to the start of an experiment to allow for metabolic recovery. For all experiments, flies were loaded into chambers by mouth pipette to avoid confounding effects of anesthesia and allowed to acclimate in the system with the air flowing for 12-24hrs prior to behavior experiments, unless otherwise specified. To control for effects of diet composition, all experimental flies were fed a consistent diet. Each chamber contained a single food vial containing 1% agar plus 5% sucrose (Sigma) with red food coloring (McCormick), which we have previously shown to result in sleep comparable to standard fly food^18^. For starvation experiments, flies had access to 1% agar dissolved in dH_2_O and were acclimated for 12 hours during lights on with access to agar alone, with analyses beginning at ZT12 at lights off. All experimental runs included analysis of both experimental flies and relevant controls in a randomized order to account for any subtle variation between runs.

### Pharmacology

Pharmacological-induced sleep was achieved through administration of gabaxodol (4,5,6,7-Tetrahydroisoxazolo[5,4-c]pyridin-3-ol hydrochloride, THIP hydrochloride; Sigma Aldrich #85118-33-8) at the dosage of 0.1 mg/ml, as previously described^19^. Gaboxadol was dissolved in dH_2_O with 1% agar and 5% sucrose. Flies were loaded into the respirometry system with the gaboxadol two hours prior to lights off (ZT10) and were maintained on the drug throughout the duration of the experiment, as described in the text.

### Sleep Deprivation

Flies were acclimated to the respirometry system during the daytime (ZT0-12). For mechanical sleep deprivation, flies were shaken every 2-3 minutes for 12 hours in the modified DAMs monitor/respirometry system throughout the night time (ZT12-24) while simultaneously measuring metabolic rate. The mechanical stimulus was applied using a vortexer (Fisher Scientific, MultiTube Vortexer) and a repeat cycle relay switch (Macromatic, TR63122). Sleep rebound and corresponding metabolic rate was measured the following day from ZT0-ZT12.

### Sleep, metabolic and statistical analyses

Respirometry recordings were analyzed using ExpeData PRO software (Sable Systems International, v1.8.4). The CO_2_ lag time from the chamber to the analyzer was corrected, the baseline was subtracted from each behavioral chamber, and the absolute CO_2_ levels (ppm) was converted to μl/hr using the recorded air flow rate. Integrating the CO_2_ trace revealed the total CO_2_ produced, or the average metabolic rate, per fly for each recording. These data were exported to Excel, where metabolic output was matched to activity, and sleep analyses were performed using a custom python program. Since individual flies were measured for either a 12hr or 24hr experimental duration (described in text), our raw data included re-sampling of metabolic rate or beam crosses for each hour. To account for these repeated hourly measures, we determined the mean of the hourly readings for each individual fly prior to our statistical analyses represented in the graphs, meaning that each fly is represented once and the “N” reported in each figure specifically refers to the number of individual flies assayed in the experiment. To detect significant differences for activity (number of beam crosses), mean VCO2 (μl/hr), or total sleep (min), we employed a student’s t-test (day vs. night; untreated control vs. gaboxadol-treated; fed vs. starved), one-way ANOVA with Sidak’s multiple comparison correction (female, day vs. night; male, day vs. night) or a two-way ANOVA with Sidak’s multiple comparison correction (*w^1118^*, fed vs. starved; *trsn*, fed vs. starved), when appropriate using InStat software (GraphPad Software 6.0). The two-tailed p-value was used to test significance is denoted as P<0.05.

To account for individual-specific differences in metabolic rate, we surveyed the metabolic rate throughout longer sleep bouts by calculating percent change in metabolic rate. This was determined by subtracting the metabolic rate during the first 5 minutes asleep from the metabolic rate during each of the subsequent 5 minutes asleep for the entire length of the sleep bout, divided by the metabolic rate during the first 5 minutes asleep, multiplied by 100 (e.g., [{first 5 min MR} – {20 min MR}/{first 5 min MR}]*100). We note some flies exhibited longer sleep bouts, however this analysis was restricted to sleep bouts up to 60 minutes due to limited replicates with extended bout lengths. Moreover, a similar approach was employed as described above for analysis of metabolic rate for flies with repeated sleep bouts. If a single fly demonstrated multiple distinct sleep bouts, we determined the mean percent change in metabolic rate for each fly at each sleep bin. For these analyses, we performed a one-way ANOVA with Sidak correction comparing the initial percent change in MR (5 min bin) to each of the subsequent sleep bins (15-60 min bins at 5 min intervals) using InStat software (GraphPad Software 6.0) with significance denoted as P<0.05.

We applied a linear regression model to characterize the relationship between both absolute vCO_2_ versus activity (number of beam crossings) and percent change in metabolic rate and sleep duration using InStat software (GraphPad Software 6.0) with significance denoted as P<0.05. Comparison of slopes derived from regression lines in fed versus starved states was performed using analysis of covariance (F-statistic; GraphPad Software 6.0). Prior to modeling, we performed pretests, including: generation of residual v. fitted plots to determine homogeneity of variance, normal Q-Q plot, Pearson Correlation table and linear model assumptions (B.L.U.E.). The culmination of these tests indicated that our data was both normally distributed and appropriate for linear regression modeling.

## Results

### Long-term recordings of sleep and metabolic rate

To simultaneously measure the effects of sleep on metabolic rate, we designed a stop-flow respirometry system coupled to a custom-built *Drosophila* Activity Monitor (DAM) system (Fig. 1A). Each DAM chamber contained three infrared (IR) beams for precise detection of locomotor activity of a single fly^20^. Humidified, fully oxygenated air was passed through each chamber, preventing desiccation and allowing for long-term recordings. After exiting the chamber, air was dehumidified and passed through a CO_2_ analyzer. The system was set to a stop-flow configuration, where the CO_2_ accumulation in each chamber was measured every 5 minutes, and these were matched to the corresponding locomotor activity of individual flies within this time period (Fig. 1B). Flies are diurnal with elevated locomotor levels during the day compared to night, and these activity patterns were maintained in the respirometry system in both male and female flies (Fig. 1C-D), indicating that the moderate airflow used in this system does not disrupt sleep-wake behavior. In both male and female flies, the mean metabolic rate was elevated during the daytime compared to the night, supporting the notion that CO_2_ production is associated with periods of high activity (Fig. 1E, F). Examination of CO_2_ levels in individual flies revealed a weak correlation in both females and males between total locomotor activity and CO_2_ levels (Fig. 1G, H). However, vCO_2_ was significantly elevated in females with activity of > 60 beam breaks and males > 50 beam breaks per five minute bin compared to the 1-10 beam breaks bin, suggesting metabolic rate is elevated during periods of robust activity (Fig. 1G, H). Therefore, this system effectively measures locomotor activity and metabolic rate simultaneously in individual *Drosophila*.

**Figure 1.**
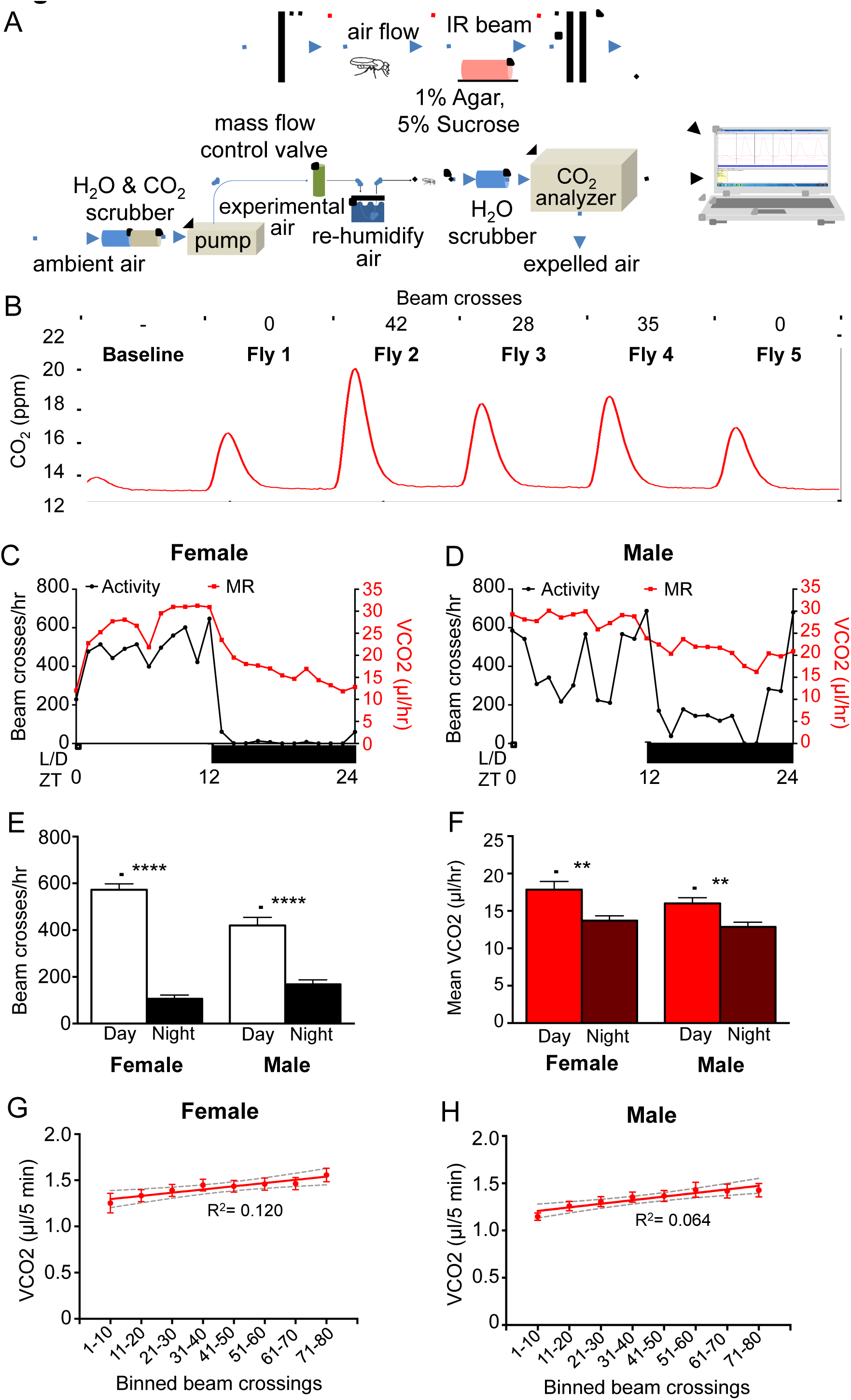
A system to measure metabolic rate in single flies. **A)** Metabolic rate was measured through indirect calorimetry. A stop-flow respirometry system measured the CO_2_ produced by single flies placed inside of a 70mm long x 20mm diameter glass tube. Each fly had access to 1% agar and 5% sucrose. Activity and sleep were measured simultaneously as metabolic rate using a *Drosophila* Locomotor Activity Monitor with three infrared beams running through each behavior chamber. The computer counted the number of beam breaks. **B)** A representative five-minute reading, with the activity in number of beam crosses and the amount of CO_2_ produced by each fly over time. **C)** The metabolic rate and activity for one female fly. **D)** The metabolic rate and activity for one male fly. **E)** The activity of female (N=24; P<0.001) and male (N=35; P<0.001) flies in beam crosses per hour, over 12 hours of day and night. **F)** The metabolic rate of female (N=24; P<0.01) and male (N=35; P<0.01) flies as CO_2_ produced per hour, over 12 hours of day and night. **G)** Linear regression of absolute vCO_2_ readout versus activity of female flies (N=24 each bin; R^2^= 0.120) and **H)** male flies (N=35 each bin; R^2^= 0.064). Grey dashed lines indicate 95% confidence interval. One-way ANOVA comparing the vCO_2_ at the 1-10 beam crossings bin to each subsequent beam crossing bin: females >60 crossings (N=24 each bin; P<0.05) and males >60 crossings (N=35 each bin; P<0.05).

### Metabolic rate is reduced in sleeping *Drosophila*

Five minutes of immobility in *Drosophila* associates with relevant behavioral and physiological sleep metrics, allowing for sleep duration to be inferred from periods of behavioral quiescence^7,10^. To measure metabolic rate during sleep, female flies were acclimated in the locomotor chambers for 24hrs, followed by continuous measurements of sleep and metabolic rate for an additional 24hrs. Flies slept significantly more during the night (ZT12-24), which corresponded with a reduction in metabolic rate (Fig. 2A-C). To determine whether changes in metabolic rate are associated with sleep bout duration, we investigated changes in CO_2_ production during sleep bouts. CO_2_ production during a single representative 60-minute sleep bout revealed a reduction in metabolic rate as sleep progressed (Fig. 2D). To account for individual variation between replicates, change in metabolic rate was calculated as percent change for each five-minute interval throughout the sleep bout compared to the first five minutes of sleep. To avoid confounds resulting from circadian differences in metabolic rate, analysis was limited to nighttime sleep. Regression analysis revealed a significant relationship between of vCO_2_ and sleep bout length (Fig. 2E). Comparing the average percent change in metabolic rate during sleep for each individual bout revealed metabolic rate was significantly reduced following 35 minutes of sleep, indicating that longer periods of uninterrupted sleep are associated with reduced metabolic rate. Percent change in metabolic rate continued to decline as sleep progressed until reaching a maximum percent change in metabolic rate of ∼-12-15% after 50 minutes of sleep. To confirm that reduction of metabolic rate during sleep is not simply due to lack of feeding activity, we compared metabolic rate during feeding and non-feeding bins from ZT1-ZT3 and did not detect significant differences in metabolic rate between feeding and waking non-feeding periods (Fig. S1). Moreover, we performed standard allometric analysis of body size versus metabolic rate to identify if weight variation among individual flies could function as a covariate affecting metabolic rate^21,22^ and determined that there is no effect of variation in body weight on metabolic rate (n=34, R^2^=0.030). Therefore, reduced CO_2_ production is associated with consolidated sleep bouts, revealing that metabolic rate can be functionally separated from overall activity.

**Figure 2.**
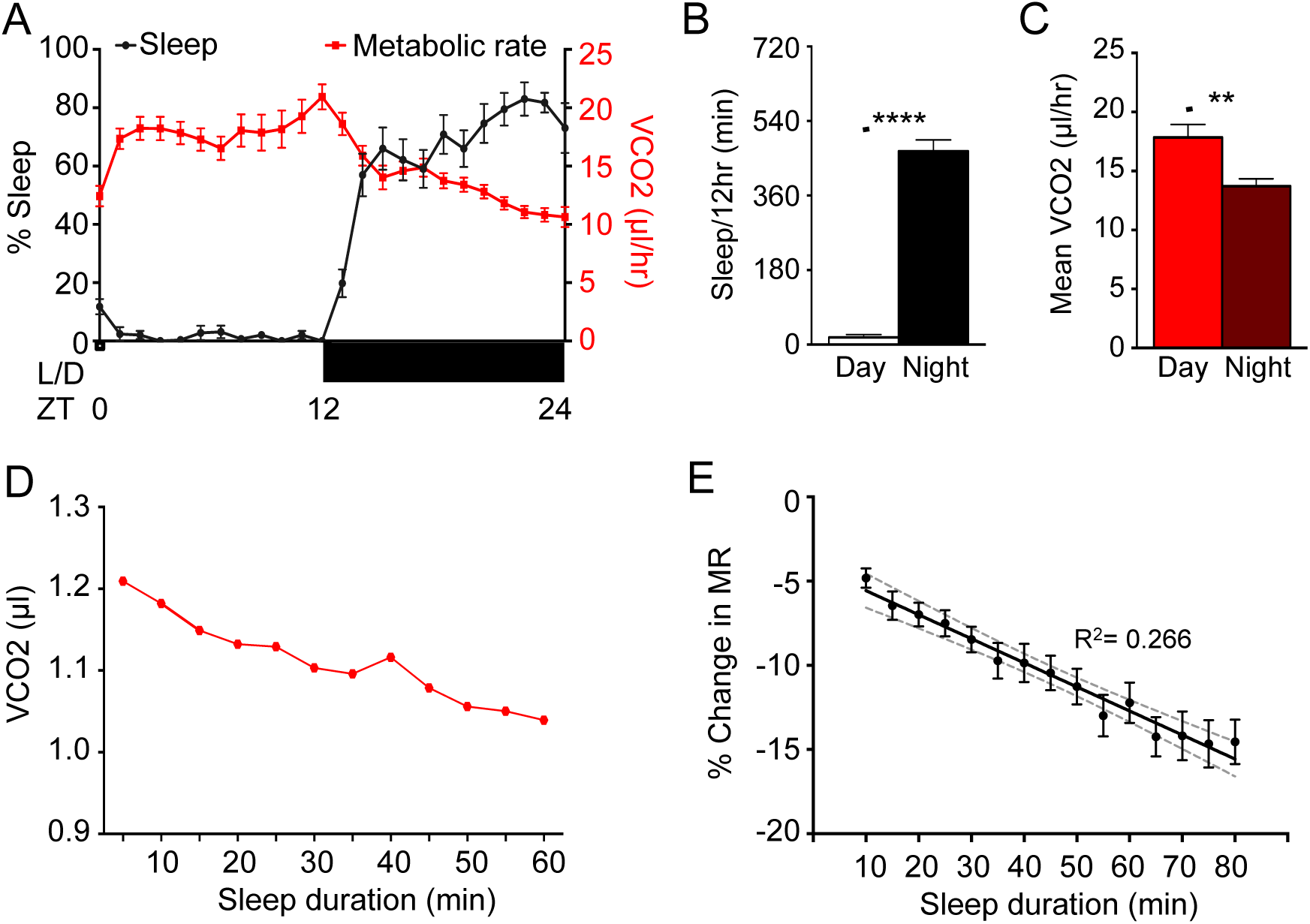
Metabolic rate is reduced during sleep state. Female control flies (*w^1118^*) were allowed to acclimate in the system for 24hrs. **A)** Metabolic rate shows an inverse pattern to their sleep (N= 24). **B)** Total minutes of sleep per 12 hours of day and night for B (N=24; P<0.001). **C)** The metabolic rate of female flies as CO_2_ produced per hour, over 24hrs of day and night (N=24; P<0.002). **D)** The metabolic rate throughout a single, representative sleep bout during the night. **E)** Linear regression model comparing percent change in metabolic rate versus sleep duration, binned per 5 minutes (N=24; R^2^=0.266). Grey dashed lines indicate 95% confidence interval. One-way ANOVA comparing the initial percent change in metabolic rate at the 10-minute sleep bin to each subsequent sleep bin reveals significant differences after 35 minutes asleep (N=24 each sleep bin; P<0.05).

### Metabolic rate during sleep deprivation and rebound sleep

To further examine the relationship between sleep and metabolic rate, we sleep deprived flies during the night (ZT12-24) and measured vCO_2_ during deprivation and recovery (Fig. 3A). Consistent with previous findings, sleep deprivation significantly increased sleep the following day (ZT0-6) compared to non-sleep deprived controls (Fig. 3B-C). Metabolic rate was elevated in sleep-deprived flies during deprivation (ZT12-24), and reduced during recovery (ZT0-6), fortifying the notion that reduced metabolic rate is associated with sleep. There was a significant correlation between metabolic rate and sleep bout duration, indicating that similar to nighttime sleep in undisturbed flies, prolonged bouts of daytime sleep are associated with reduced metabolic rate (Fig. 3F). Rebound sleep demonstrated a significant reduction in metabolic as sleep duration progressed beyond 35 minutes (Fig. 3F), further supporting the notion that daytime rebound recapitulates physiologically similar sleep-associated metabolic changes to nighttime sleep.

**Figure 3.**
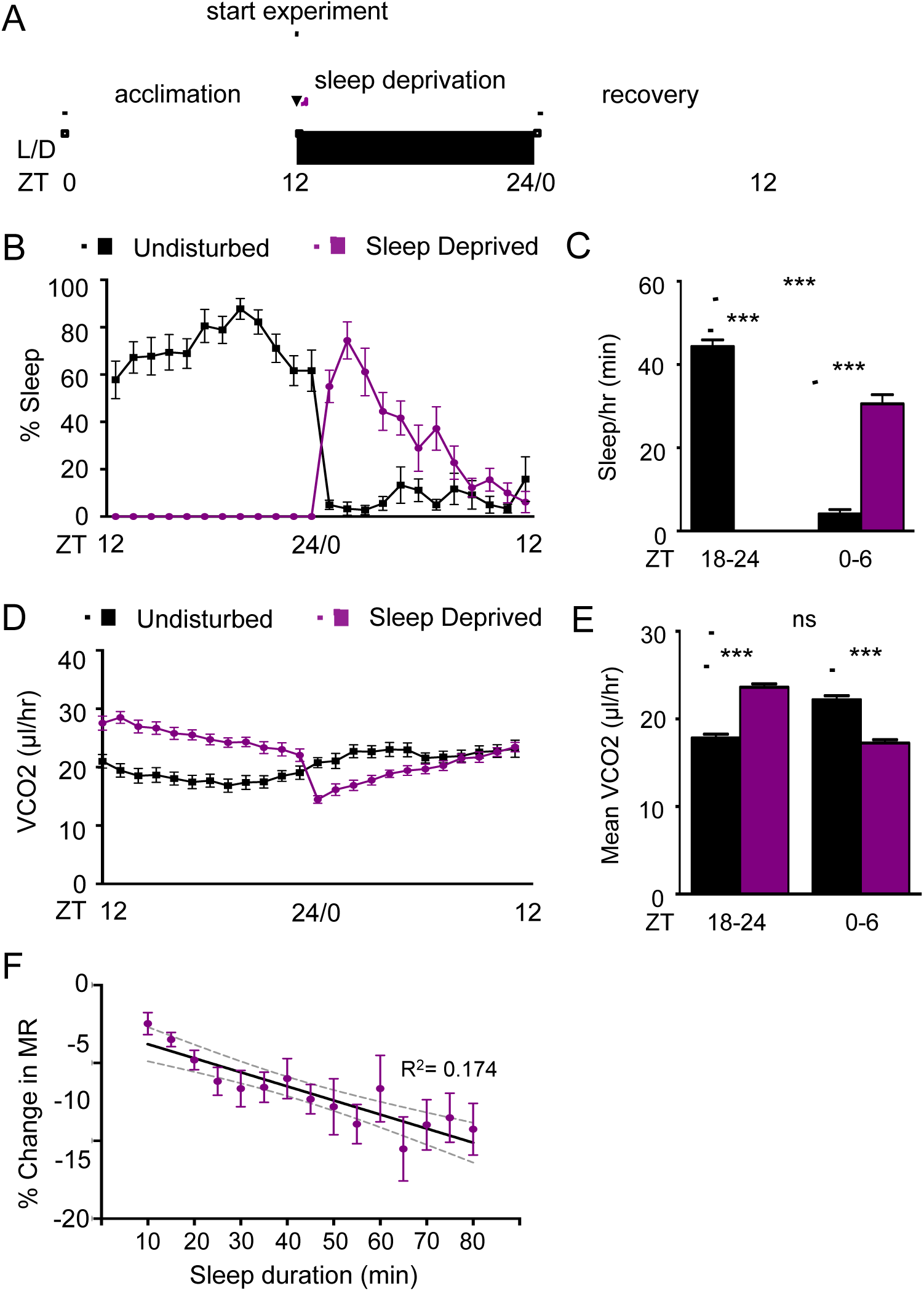
Metabolic rate is elevated during sleep deprivation and reduced during rebound. **A)** Female control flies (*w^1118^*) were acclimated during the day (ZT0-12). Mechanical sleep deprivation was applied during the 12 hr night (ZT12-24) and recovery was assessed the following day (ZT0-12). **B)** Sleep deprived flies (N=15; **purple)** sleep more during the first 6 hours of daytime following deprivation (ZT0-6) relative to undisturbed controls (N=15; black). **C)** Quantification of total sleep shows that flies were sufficiently sleep deprived during nighttime (ZT18-24; P<0.0001) and demonstrated increased sleep during the recovery period (ZT0-6; P<0.0001). **D)** Hourly profile of metabolic rate in sleep deprived and control flies. **E)** Quantification metabolic rates demonstrates elevated metabolic rate during sleep deprivation (ZT18-24; P<0.0001) and reduced metabolic rate during recovery (ZT0-6; P<0.0001). Metabolic rate during recovery in sleep deprived flies is comparable to levels of control flies during normal nighttime sleep (P>0.327). **F)** Regression analysis comparing percent change in metabolic rate versus sleep duration, binned per 5 minutes (N=15; R^2^= 0.174). Grey dashed lines indicate 95% confidence interval. One-way ANOVA comparing the initial percent change in metabolic rate at 10- minute sleep bin to each subsequent sleep bin reveals significant differences after 35 minutes asleep (N=15 each sleep bin; P<0.05).

### Metabolic rate is reduced during pharmacologically-induced sleep

GABA signaling promotes sleep in diverse species and the GABA-A Receptor agonist gabaxodol potently induces sleep in *Drosophila*^19,23–25^ To determine the effects of pharmacologically-induced sleep on metabolic rate, we housed flies on agar containing 0.1 mg/ml gaboxadol and 5% sucrose in the respirometry chambers and measured the effects on sleep and metabolic rate (Fig. 4A). Consistent with previous studies, sleep was elevated in gaboxadol-treated flies compared to controls throughout the 12-hour daytime recording^19^ (Fig. 4B,C). Notably, metabolic rate was reduced in gaboxadol-treated flies during the daytime compared to controls, confirming that pharmacologically induced sleep lowers metabolic rate (Fig. 4D, E). These experiments were limited to analysis of daytime sleep, therefore, we could not determine percent change in metabolic rate of untreated *w^1118^* flies across sleep bouts, since control flies sleep very little during the day in this paradigm. Sleep bout length in gaboxadol-treated flies was associated with reduced metabolic rate (Fig. 4F). Moreover, comparison of the of percent change in metabolic rate of each subsequent sleep bin relative to the first change at 10 minutes show a robust reduction in metabolic rate in gaboxadol-treated flies after 30 minutes of sleep (Fig. 4F). Because the percent change in metabolic rate is comparable to the metabolic rates of wild-type flies during night sleep, it is possible that pharmacologically-induced daytime sleep is physiologically comparable to nighttime sleep.

**Figure 4.**
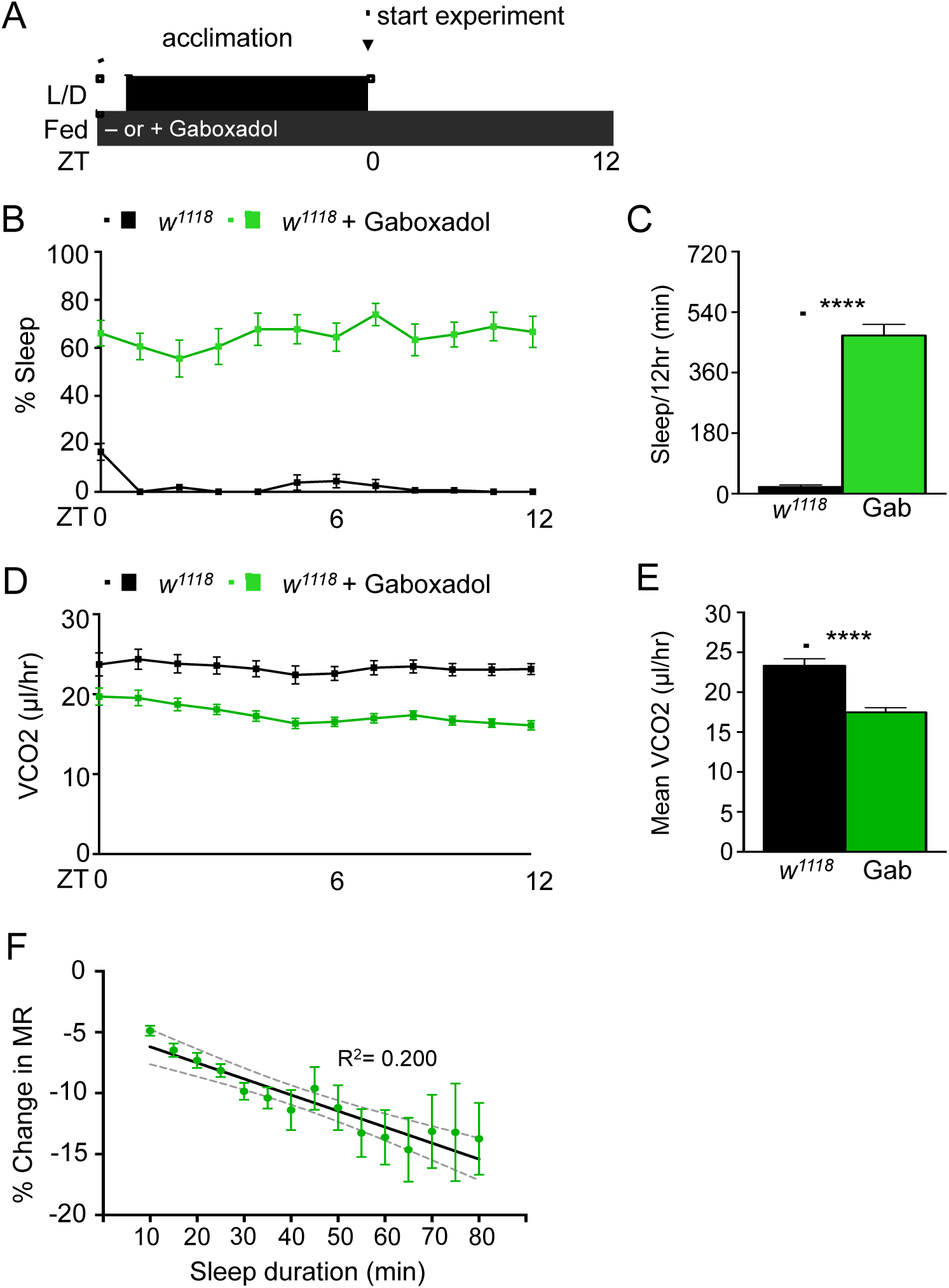
Reduced metabolic rate during pharmacologically-induced sleep. **A)** Female *w^1118^* flies were loaded on sucrose or sucrose containing 0.1 mg/ml gaboxadol two hours prior to lights out (ZT10), acclimated to the system for 12 hours during the night phase and were measured for 12 hours (ZT0-12) during the following day. **B)** Daytime sleep was significantly elevated in gaboxadol-treated flies (green) compared to flies fed sucrose alone (black). **C)** Quantification of total sleep reveals gaboxadol-treated flies (N=15) sleep significantly longer than untreated controls (N=14; P<0.0001). **D)** Metabolic rate was reduced throughout the 12 hour day. **E)** Quantification of mean metabolic rate reveals a significant reduction in gaboxadol-treated flies (N=15) compared to controls (N=14; P<0.0001). **F)** Linear regression of percent change in metabolic rate versus sleep duration, binned per 5 minutes (N=15; R^2^= 0.200). Grey dashed lines indicate 95% confidence interval. One-way ANOVA comparing the initial percent change in metabolic rate at the 10-minute sleep bin to each subsequent sleep bin reveals significant differences after 30 minutes asleep. (N=15 each sleep bin; P<0.05).

### The effects of starvation on sleep and metabolic rate

In mammals, starvation potently suppresses sleep and metabolic rate^26^. Further, flies suppress sleep shortly after the onset of food deprivation, presumably to increase foraging behavior^18,27^. To determine how starvation-induced sleep suppression impacts metabolic function, we compared the metabolic rate of fed and starved female *w^1118^* flies. Flies were acclimated for 12 hours on food or agar, and metabolic rate was measured during the 12 hour night phase (ZT12-24; Fig. 5A). In agreement with previous findings, sleep was reduced in starved flies throughout the 12hr recording period^28,29^ (Fig. 5B, C). Despite the loss of sleep in starved flies, metabolic rate was lower in starved animals compared to fed counterparts, providing further support that metabolic rate in *Drosophila* is modulated independently from locomotor activity (Fig. 5D, E). There was a significantly stronger relationship between sleep bout length and metabolic rate in fed flies, providing evidence that starvation impairs sleep-associated physiological changes on metabolic rate (Fig. 5F). To determine the effect of starvation on sleep-dependent regulation of metabolic rate, we compared the metabolic rate of each sleep bout in fed and starved animals. In fed flies, metabolic rate was reduced following 40 minutes of sleep compared to the first 5 minutes of sleep, yet when starved, metabolic rate is not significantly reduced as sleep progresses, further supporting the notion that starvation impedes sleep (Fig. 5F). Taken together, these findings reveal that CO_2_ production is reduced in starved flies without affecting sleep-dependent changes in metabolic rate.

**Figure 5.**
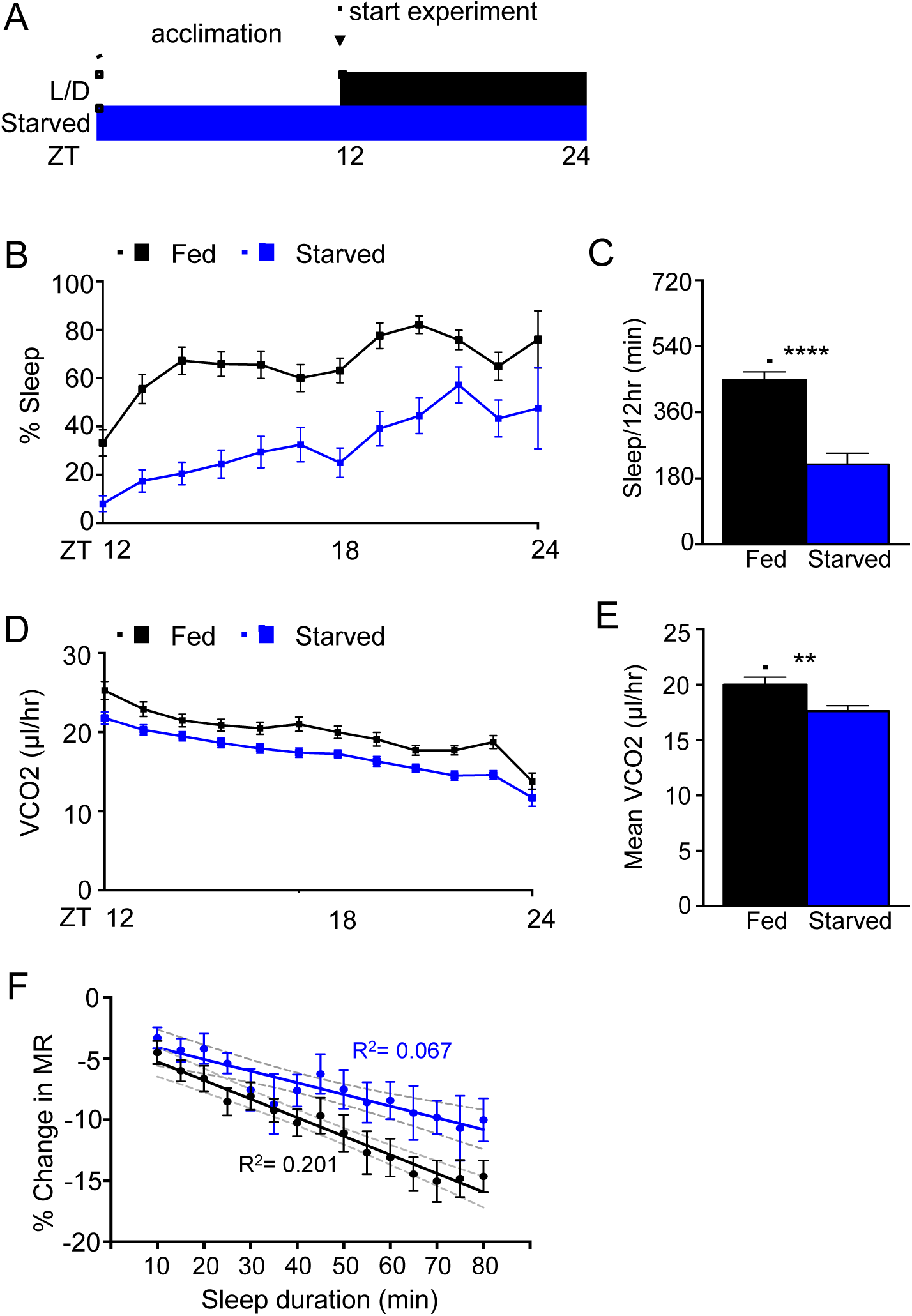
Metabolic rate and sleep are reduced in starved flies. **A)** Flies were fed or starved while acclimating to the system for 12 hours during the day (ZT0-12) prior to measurement throughout the night (ZT12-24). **B)** Flies starved on agar (blue) slept less than flies housed on 5% sucrose (black) during the 12 hour night period. **C)** Quantification of total sleep over the 12 hour night period reveals a significant reduction in starved flies (N=30) compared to fed controls (N=29; P< 0.0001). **D)** Metabolic rate is lower throughout the 12hr nighttime period in starved flies. **E)** Quantification of mean vCO_2_ production over this period reveals a signification reduction in starved animals (N=30) relative to controls (N=29; P<0.01). **F)** Regression analysis comparing percent change in metabolic rate versus sleep duration, binned per 5 minutes reveals a correlation in fed flies (N=29; R^2^= 0.201), but little effect in starved flies (N=26 each sleep bin, 4 flies did not have any sleep bouts when starved; R^2^=0.067). Grey dashed lines indicate 95% confidence interval of each line. Comparison of the regression lines indicate that the slopes are different between the fed versus starved state (F=5.319; P<0.05). One-way ANOVA comparing the initial percent change in metabolic rate at the 10-minute sleep bin to each subsequent sleep bin within each group reveals significant differences after 40 minutes asleep in fed flies (N=29 each sleep bin; P<0.05) and no significant differences in starved flies (N=26 each sleep bin).

### Metabolic changes during sleep are intact in *translin* mutant flies

It is possible that shared genes regulate starvation-induced reductions in sleep duration and sleep-dependent regulation of metabolic rate. We previously identified the RNA binding protein *translin* (*trsn*), as essential for starvation-induced sleep suppression^29^. Energy stores and feeding behavior are normal in *trsn* deficient flies, yet they fail to suppress sleep in response to starvation, suggesting *trsn* is required for the integration of sleep and metabolic state^29^. To determine whether *trsn* affects metabolic rate, we measured sleep and metabolic rate in fed and starved *trsn* mutant flies. Flies were loaded into the respirometry system and allowed to acclimate for 12hrs during the day. Sleep and metabolic rate were then measured for the duration of the night phase (ZT12-24). In agreement with previous findings, control flies robustly suppressed sleep when starved on agar, while there was no significant effect of starvation on sleep duration in trsn^null^ flies (Fig. 6A, B). In both control and trsn^null^ flies, CO_2_ production was reduced during starvation, suggesting *trsn* is not required for modulating metabolic rate in accordance with feeding state. These findings fortify the notion that metabolic rate can be regulated independently from both sleep and locomotor activity (Fig. 6C, D). Interestingly, while metabolic rate was further reduced in *trsn*^null^ flies upon starvation, the basal metabolic rate of fed *trsn^null^* flies was lower than *w*^*1118*^ controls (Fig. 6C, D). For both *w^1118^* and *trsn*^null^ flies, there was a stronger relationship between metabolic rate and sleep bout duration in fed flies than starved flies (Fig. 6E, F), fortifying the notion that sleep-dependent changes in metabolic rate are not disrupted in *trsn*^null^ flies. Together, these results indicate that *trsn* is required for starvation-induced sleep suppression, but is dispensable for sleep-induced modulation of metabolic rate.

**Figure 6.**
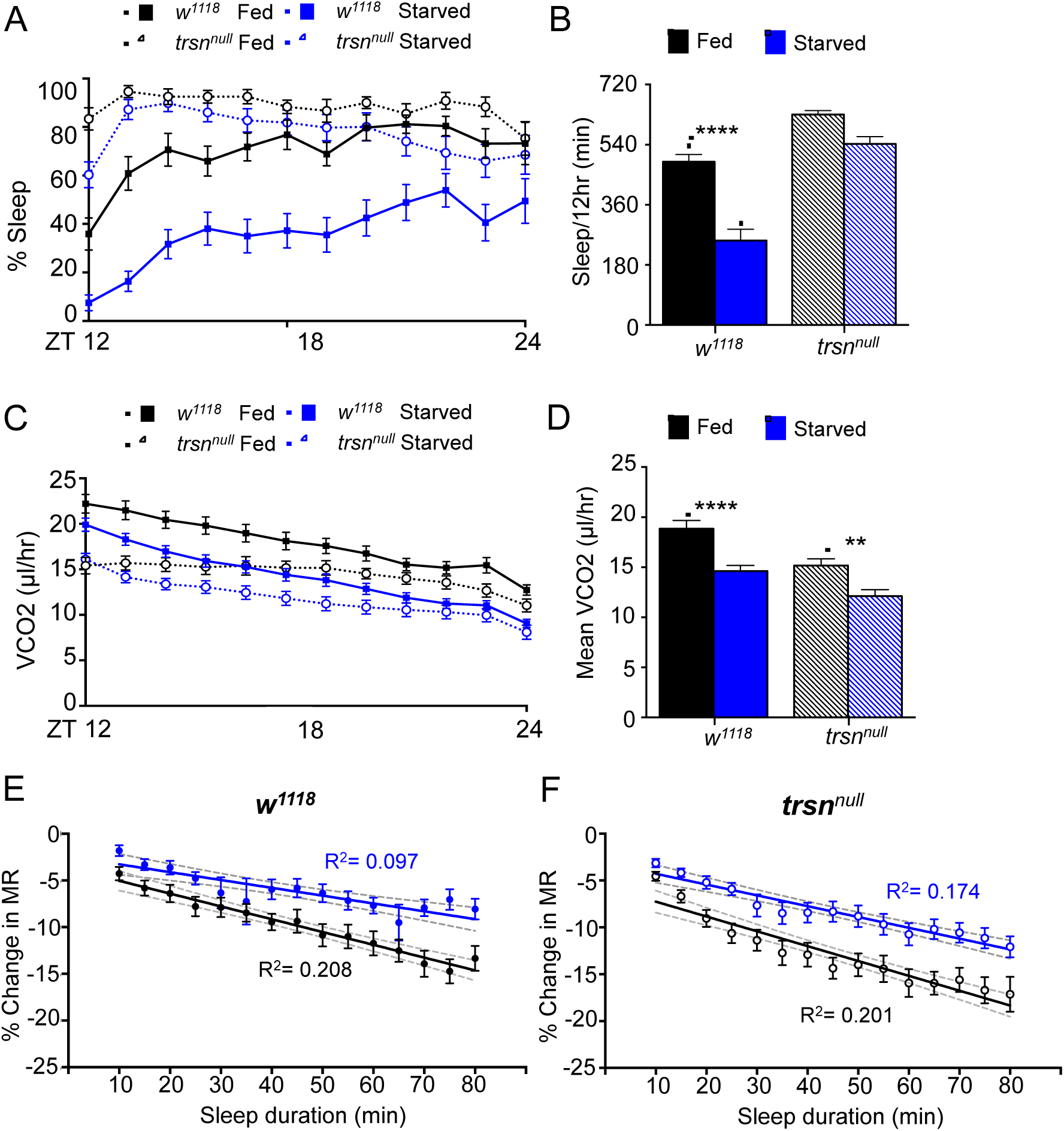
Sleep-dependent changes in metabolism are intact in *trsn^null^* flies. **A)** Sleep did not significantly differ between *trsn^null^* flies housed on sucrose or starved on agar alone. **B)** Quantification of total nighttime sleep (ZT12-ZT24) revealed sleep is significantly lower in *w^1118^* control flies (N=30) housed on agar compared to fed (N= 28; P<0.0001), while there is no significant difference between *trsn*^null^ flies (N=28) housed on 5% sucrose or agar alone (N= 25; P>0.05). **C)** Metabolic rate is lower in control and *trsn*^null^ flies housed on agar compared to flies housed on 5% sucrose. **D)** Quantification revealed metabolic rate is lower in both *trsn*^null^ flies and controls (*w^1118^*, P<0.0001; *trsn^null^*, P<0.01). **E)** Applied linear regression model comparing percent change in metabolic rate versus sleep duration, binned per 5 minutes reveals a correlation in *w^1118^* fed flies (N=28; R^2^= 0.208), but only a weak effect in *w^1118^* starved flies (N=25, 5 flies did not sleep on agar; R^2^=0.097). Grey dashed lines indicate 95% confidence interval of each line. Comparison of the regression lines indicate that the slopes are different between the *w^1118^* fed versus starved state (F= 7.09725, P<0.01). One-way ANOVA comparing the initial percent change in metabolic rate at the 10-minute sleep bin to each subsequent sleep bin within each group reveals significant differences after 40 minutes asleep in fed flies (N=28 each sleep bin; P<0.05) and differences in starved flies beyond 55 minutes (N=25 each sleep bin; P<0.05). **F)** Regression analysis model comparing percent change in metabolic rate versus sleep duration, binned per 5 minutes reveals a correlation in *trsn*^null^ fed flies (N=28; R^2^= 0.201), but only a weak effect in *trsn*^null^ starved flies (N=25; R^2^=0.183). Grey dashed lines indicate 95% confidence interval of each line. Comparison of the regression lines indicates that the slopes do not differ between the *trsn*^null^ fed versus starved state (F=5.0557, P<0.05). One-way ANOVA comparing the initial percent change in metabolic rate at the 10-minute sleep bin to each subsequent sleep bin within each group reveals significant differences after 25 minutes asleep in trsn^null^ fed flies (N=28 each sleep bin; P<0.05) and differences in *trsn*^null^ starved flies beyond 30 minutes (N=25 each sleep bin; P<0.05).

## Discussion

Metabolic rate is regulated in accordance with environmental changes and life history, providing a metric for whole-body metabolic function. While mammalian studies typically determine metabolic rate via O_2_ consumption or respiratory quotient (ratio of CO_2 eliminated_/O_2 consumed_), studies in *Drosophila* commonly measure CO_2_ production because it is directly correlated with O_2_ input and accurately reflects metabolic rate^14,30,16^. Previous systems investigating metabolic rate in *Drosophila* have used single flies or populations to measure changes in CO_2_ in response to aging, temperature change and dietary restriction^16,31,32^. Here, we have modified a previously described single-fly respirometry system and *Drosophila* activity monitor system to simultaneously measure metabolic rate and sleep. This system can measure CO_2_ production and locomotor activity over a 24hr period, providing the ability to measure the relationship between sleep and metabolism, providing a system to investigate the complex relationships between diverse genetic and environmental factors with metabolic rate.

### Sleep-metabolism interactions in mammals and arthropods

Regulation of sleep and metabolism are conserved at the molecular and physiological levels between *Drosophila* and mammals^33,34^. Similar to mammals, flies modulate sleep and feeding in accordance with metabolic state, providing a system to investigate the genetic underpinnings of these behaviors^2^. For instance, when starved, both flies and mammals suppress sleep presumably to forage for food^18,35,36^. The finding that sleep-dependent reductions in metabolic rate are conserved in *Drosophila* supports the notion that an essential function of sleep is metabolic regulation. A number of previous studies suggest total sleep duration is positively correlated with basal metabolic rate, supporting the notion that sleep may be an adaptive mechanism of energy conservation^37,38^. However, a meta-analysis study examining over 40 different mammalian species revealed a negative relationship between sleep and basal metabolic rate, opposing the energy conservation model of sleep^39^. In humans, reduced metabolic rate during sleep accounts for as much as a 15% energy savings^40,41,42^. Our findings reveal a similar reduction of metabolic rate during sleep in fruit flies, suggesting this may provide an evolutionarily adaptive mechanism to conserve energy.

While our study is the first to examine the relationship between sleep and metabolic rate in *Drosophila*, previous studies implicated shared genetic or environmental factors in the regulation of sleep and metabolic function. For example, dopamine potently suppresses sleep in *Drosophila*, and flies harboring a mutation in the dopamine transporter gene *fumin* (*fmn*) exhibit reduced sleep and increased CO_2_ production^43,44^, suggesting dopamine regulates both sleep and metabolic state. Importantly, metabolic rate remained elevated in *fumin* mutant flies when motor neurons were genetically silenced, indicating that the elevated metabolic rate does not result from differences in locomotor activity^44^. Moreover, long-term sleep deprivation in Pacific beetle cockroach, *Diploptera punctate*, caused significant increases in O_2_ consumption and elevated basal metabolic rate compared to controls, indicating that sleep loss impacts metabolism^45^. These data are in agreement with our findings, where we identify increased metabolic rate during sleep deprivation and reduced metabolic rate during sleep rebound or pharmacologically-induced sleep, revealing a fundamental, and direct relationship between sleep and lower metabolic rate, indicating that changes in CO_2_ production during sleep are due to changes in basal metabolic rate, rather than reduced locomotor activity.

### Environmental factors regulating sleep and metabolism

In *Drosophila*, sleep and metabolic rate are influenced by diet, temperature and age^46,47^. Here, we discover that starvation conditions impede the physiological changes associated with normal sleep. This simultaneous assessment of sleep and metabolic state can be applied to determine how metabolic rate and sleep are related to starvation resistance. Selection for starvation-resistant *Drosophila* through experimental evolution results in flies that can survive over two weeks without food and exhibit a host of metabolic and developmental differences, thus providing a system to examine interactions between metabolism and behavior^48,49^. The starvation-resistant flies exhibit increased body size, energy stores and reduced metabolic rate, providing numerous mechanisms for energy conservation^49–51^. Previously, we reported that sleep duration is increased in starvation resistant flies, and proposed that this provides an additional mechanism for energy conservation^50,52^. Application of this approach measuring sleep and metabolic rate will provide the ability to determine metabolic rate in asleep and awake flies and identify whether reduced metabolic rate in starvation-resistant flies is a consequence of increased sleep, or these traits have evolved in parallel.

### Evidence for sleep-associated regulation of metabolic rates in *Drosophila*

In birds and mammals, sleep is associated with changes in cortical activity resulting in defined stages of sleep, such as REM and NREM, which differ in physiology and function^53,54^. In *Drosophila*, sleep studies have primarily used behavioral quiescence and body postures to denote sleep, and much less is known about how sleep impacts physiology. Recording of local field potentials in tethered animals reveals distinct differences between quiescent and active states, and sleep is associated with a reduction in 15-30Hz local field potentials. The reduction in 15-30Hz oscillations is at its greatest following 15 minutes of immobility, suggesting that this physiological change in neuronal activity represents a deeper form of sleep, along with coordinate increases in arousal threshold^9,10^. Evaluating sleep intensity in male and female *Drosophila* using an arousal-testing paradigm during extended nighttime sleep bouts identified a gradual decrease in responsiveness until a second, deeper sleep state was reached after ∼30 minutes^10,55^. Consistent with these findings, we report that metabolic rate decreases with sleep duration, reaching a significant reduction ∼30 minutes following sleep onset. Taken together, these findings suggest metabolic rate may provide a physiological indicator of sleep intensity that compliments existing electrophysiology and behavioral methods of analysis to define a deeper sleep state in flies.

### A role for *translin* in regulating metabolic rate

The RNA binding protein *trsn* is a proposed integrator of sleep and metabolic state, and flies deficient for *trsn* fail to suppress sleep in response to starvation^29^. Notably, the defect in *trsn* mutant flies is specific to regulation sleep regulation, because *trsn* deficient flies have normal feeding and energy stores^29^. Here, we find that starvation reduces metabolic rate in *trsn* mutant and wild-type flies. Even though *trsn* mutants demonstrate starvation-induced sleep suppression, our findings indicate that metabolic rate can still be modulated in *trsn* mutants in a starved state. We identify a lower basal metabolic rate in fed *trsn* mutants compared to controls, though this may be attributable to the trend towards increased nighttime sleep in *trsn* mutant flies. In addition to *trsn*, a number of additional genes and transmitters have been identified as regulating starvation-induced sleep suppression or hyperactivity, including Octopamine, *clock*, and the glucagon-like *adipokinetic* hormone^35,56,57^. Therefore, this assay provides a direct readout of metabolic response to starvation and can be used for more detailed investigation of the mechanisms underlying the integration of sleep and metabolic state.

### Future applications for investigation of metabolic rate in *Drosophila*

Beyond our initial analysis of the relationship between sleep and metabolic rate in *Drosophila*, this system allows for genetic screens or targeted genetic manipulations to identify novel genes regulating sleep, metabolic rate, and the integration of these processes. For example, the mushroom bodies, fan-shaped body, and circadian neurons modulate sleep and wakefulness in *Drosophila*^23,58–61^ and the effects of manipulating these systems on sleep-dependent modulation of metabolic rate could be measured using this system. Similarly, application of this system could identify novel genes, neurons or environmental factors required for changes in metabolic rate during sleep. Recent studies in flies have identified neural circuits involved in sleep homeostasis^62–64^, yet little is known about the physiological changes associated with rebound sleep in flies. In addition to sleep, the circadian system regulates metabolism in flies and mammals^65,66^.

In addition to measuring metabolic rate, respirometry can also be used to measure specific molecules being metabolized. The simultaneous measurement of CO_2_ and O_2_ enables the ability to identify the exchange ratios of O_2_ consumed and CO_2_ produced, and ultimately infer the specific energy fuels utilized^67,68^. More specifically, the ratio between the CO_2_ produced and O_2_ consumed at a steady state, also known as the respirometry quotient (RQ), or the respirometry exchange ratio (RER) which quantifies the same ratio but at any time point (e.g., during exercise), can be used to identify food sources metabolized including fat (RQ= ∼0.7), carbohydrates (RQ= ∼1.0) or protein (RQ= ∼0.8-0.9)^14^. Respirometry measurements have been used to identify substrates metabolized in both mammals^69,70^ and invertebrates^16^. However, the sensitivity of O_2_ detection is lower than CO_2_, thus preventing detection of O_2_ changes in single flies, yet recent studies indicate that sleep can be measured in group-housed *Drosophila*^71^. Therefore, it may be feasible for future studies utilizing groups of flies to determine metabolized energy stores using this system.

## Conclusions

We describe a system for simultaneously measuring sleep and metabolic rate, and further, identify dynamic regulation of metabolic rate during individual sleep bouts. This system denotes metabolic rate as a readily identifiable marker of the physiological changes associated with sleep, which can be universally applied to examine the function of novel sleep genes and neurons in *Drosophila*. Ultimately, this unique system can be applied to examine precise interactions between numerous aspects of life history and circadian function coordinately with metabolic rate.

## Acknowlegements

This work was supported by NIH grants 1R01NS085252 to ACK and R15NS080155 to ACK and JRD. The authors are grateful to Dr. Allen Gibbs (University of Nevada, Las Vegas) for initial guidance in establishing the respirometry system, as well as Mark Spencer (Trikinetics) and Dr. Thomas Foerster (Sable Systems), for technical advice and assistance designing this system. Dr. Paul Shaw (Washington University, St. Louis) provided critical experimental suggestions and feedback.

---

**Figure S1. Metabolic rate does not differ in active flies during feeding versus nonfeeding bins. A)** Female flies were mouth pipetted individually into glass behavioral chambers the night before experiment start to allow for at least 12 hours of acclimation. Flies were then tested at approximately ZT1 or ZT3 for simultaneous measures of metabolic rate, activity, and feeding through video recording fly behavior. Feeding behavior was manually scored, and was qualified as proboscis extension to the food source. The respirometry system was flushed of residual air for 30 minutes prior to experiment start. Sampling alternated between the fly and a baseline chamber every minute, to give the total VCO_2_ (μl) produced by the fly every 2 minutes. Experiment duration was 1 hour. **B)** Representative metabolic rate, activity, and feeding results for one fly. Metabolic rate does not fluctuate dramatically between feeding and non-feeding periods. **C)** Mean metabolic rates did not significantly differ between feeding and non-feeding periods (P>0.380). Mean metabolic rates during both feeding and active states were significantly higher than the mean sleep metabolic rate (N=16 flies). Feeding v sleeping (P<0.01) and active versus inactive (P<0.05).

